# Combining Crowdsourcing and Deep Learning to Assess Public Opinion on CRISPR-Cas9

**DOI:** 10.1101/802454

**Authors:** Martin Müller, Manuel Schneider, Marcel Salathé, Effy Vayena

## Abstract

The discovery of the CRISPR-Cas9-based gene editing method has opened unprecedented new potential for biological and medical engineering, sparking a growing public debate on both the potential and dangers of CRISPR applications. Given the speed of technology development, and the almost instantaneous global spread of news, it’s important to follow evolving debates without much delay and in sufficient detail, as certain events may have a major long-term impact on public opinion and later influence policy decisions. Social media networks such as Twitter have shown to be major drivers of news dissemination and public discourse. They provide a vast amount of semi-structured data in almost real-time and give direct access to the content of the conversations. Such data can now be mined and analyzed quickly because of recent developments in machine learning and natural language processing. Here, we used BERT, an attention-based transformer model, in combination with statistical methods to analyse the entirety of all tweets ever published on CRISPR since the publication of the first gene editing application in 2013. We show that the mean sentiment of tweets was initially very positive, but began to decrease over time, and that this decline was driven by rare peaks of strong negative sentiments. Due to the high temporal resolution of the data, we were able to associate these peaks with specific events, and to observe how trending topics changed over time. Overall, this type of analysis can provide valuable and complementary insights into ongoing public debates, extending the traditional empirical bioethics toolset.

## 1 Introduction

Genome editing has many potential applications, ranging from gene therapy [1] to crop enhancement [2] and production of biomolecules [3, 4]. While it has been possible to modify the genomes of eukaryotic cells since the 1980s, traditional methods have proven to be rather impractical, inaccurate or impossible to use at scale [5, 6, 7, 8]. Accurately targeted gene editing has only become possible within the last decade [9, 10] using a CRISPR-Cas9 based method. In 2013, the method was further developed to be used on human cells [11, 12], which allowed for the first successful experiment to alter the human germline DNA of non-viable embryos in April 2015 [13]. The experiment, conducted by a group of Chinese scientists, raised ethical concerns among researchers and the general public about the potential far-reaching consequences of introducing germline modifications [14, 15]. Such ethical concerns include unexpected side effects on the evolution of humans, as well as cultural and religious arguments. In November 2018, Jiankui He announced the genetic editing of two viable human embryos with the goal of introducing HIV resistance [16]. The work came to be known to a global public under the term “CRISPR babies”, and was condemned by the scientific community as unethical, unnecessary and harmful to the two babies [17, 18].

As the costs of the technology drop further and usage becomes more widespread, governments and policy makers are faced with the challenging task of posing adequate ethical restrictions to prevent misuse. In order to gain time to introduce appropriate ethical frameworks, some scientists have called for a moratorium on genetically editing the human germline [19, 20, 21]. Previous studies on opinion towards GMO plants highlight how certain events or scandals (e.g. with respect to food safety) may have a major long-term impact on public opinion and later drive policy decisions [22, 23, 24, 25]. Understanding the public attitudes towards topics such as CRISPR is therefore of paramount importance for policy making [26, 27].

Several surveys have been conducted with the goal of evaluating the public’s perception of CRISPR and genetic engineering in general [28, 29, 30, 31, 32]. Such surveys have found that participants are largely in favor of the technology used for somatic purposes (e.g. in the context of treatment) but less so for germline editing, especially if this is not for clearly medical purposes. Additionally, the studies underline certain demographic correlations, e.g. that women, people belonging to ethnic minorities, and religious communities are more critical about the potential applications of CRISPR [30, 28]. Somewhat unsurprisingly, the surveys also show that public views are not always aligned with expert opinions [32]. A recent study that explored coverage of news articles on CRISPR in North America between 2012 and 2017, found CRISPR to be overwhelmingly portrayed as positive and potentially overhyped in news media compared to the public’s views [33].

In this study, we provide the first analysis of a complete dataset of all tweets about CRISPR published over a six and a half year period. The analyzed timespan includes the first experiment of CRISPR on human cells in 2013 but also recent events, such as the first genetic editing of viable human babies in November 2018. Furthermore, we make use of recent advances in text classification models, such as BERT [34], which use semi-supervised machine learning to generate a high-resolution temporal signal of the sentiment towards CRISPR over the observed timespan. By combining multiple text classification methods, we obtain results which can also be linked back to previous studies conducted with traditional methods, such as surveys.

## 2 Methods

Our analysis consists of four different, explorative approaches, all of which build upon the sentiments of the tweets. Therefore, sentiment analysis represents the core of our analysis. In order to determine the sentiment for the entirety of tweets published over the last six and a half years, we trained a predictive model on a previously manually annotated subset of the data. The process can be divided into five main tasks: Data collection, preparation, annotation, training, and analysis, which we describe in the following (see Figure 1 for an overview of the process).

**Figure 1:**
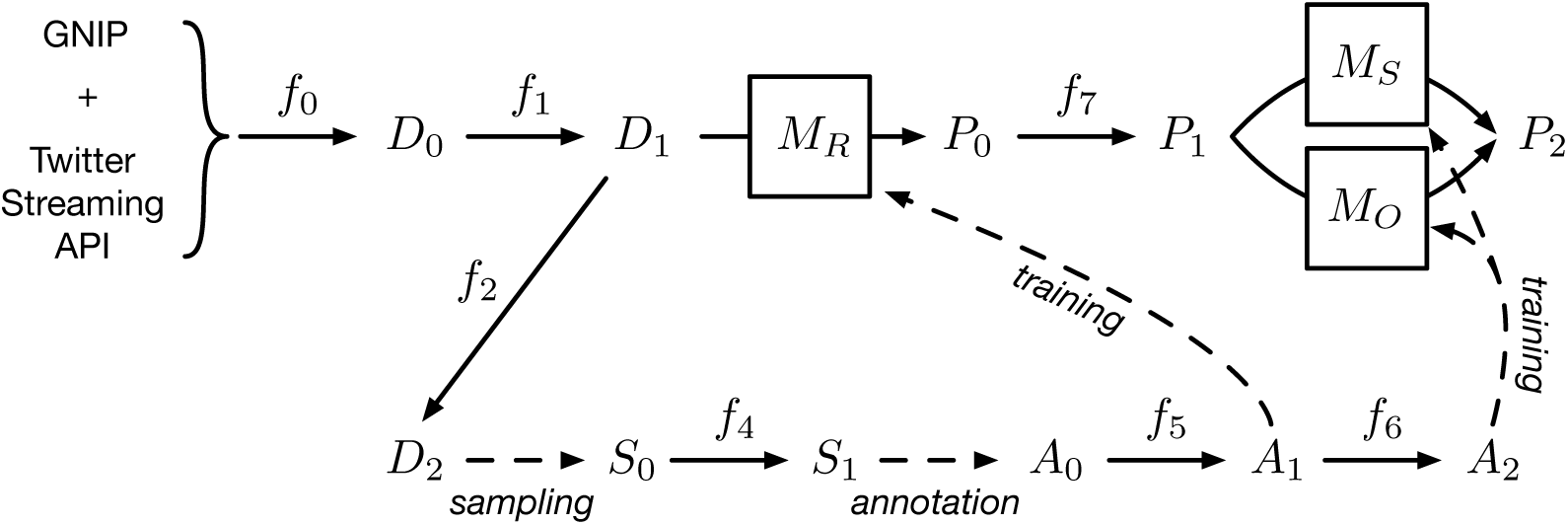
Overview of the data processing pipeline. Raw data is preprocessed by multiple filtering steps *f*_0−2_ into different forms, denoted as *D*_0−2_. From these data sets, a sample *S*_0_ is drawn and subsequently annotated into *A*_0−2_. Finally, three models (*M*_*R*_, *M*_*S*_, and *M*_*O*_) are trained on the annotated data and used to predict the data set *D*_1_, resulting in predicted data *P*_0−2_. Please refer to the text for details and a description of all filtering steps *f*_0−7_.

### 2.1 Data collection

The data set (denoted as *D*_0_ in Figure 1) for our analysis consists of all tweets (including retweets, quoted tweets, replies, and mentions) that match the character sequence CRISPR (in any capitalization), have been detected to be in English language and were published between January 1, 2013 and May 31, 2019. We retrieved this data either through the Twitter Streaming API or through GNIP, a Twitter subsidiary which allows access to historical data which was not retrievable through the Twitter Streaming API. The three filtering conditions mentioned above were used as parameters in the retrieval through Twitter APIs (denoted as *f*_0_) as well as for the requested data from GNIP.

The number of tweets varies greatly over time, ranging from 4818 in 2013 up to 445 723 in 2018, totaling 1 508 044 tweets by 348 502 distinct users (also refer to Table A.1). Since the focus lies on the overall evolution of the discourse provided by aggregated information, the study considers only the text in the tweet objects and ignores user-related information (such as location) or media content (such as photos or videos). In addition, any occurrences of Twitter handles and URLs in the text were anonymized (replaced by @<user> and `<url>`) to protect individuals.

### 2.2 Preparation

In a preparatory step, tweets suitable for annotation were selected from *D*_0_. As an inclusion criterion, only tweets with at least three English words (after removal of stop words) were considered (*f*_1_).

Although a tweet with less than three non-stop-words may express a sentiment, we chose this threshold to ensure that the annotators had at least a minimal context to determine if the tweet was in fact relevant to the topic, and what sentiment it expressed. The word count was determined by the help of NLTK’s TweetTokenizer and English word and stop word corpora [35]. The resulting dataset *D*_1_ (*n* = 1 334 114) was used as the basis for the subsequent analysis. In order to avoid the annotation of duplicates, all retweets, quoted tweets, and other duplicates of tweets with the same text were removed, leading to dataset *D*_2_ (*n* = 433 930).

Next, we selected a random sample *S*_0_ (*n* = 29 238), so that we obtained a more or less evenly distributed number of tweets over the observed timespan. This was achieved by binning the data by all 77 months and selecting a constant number of tweets from each monthly bin. In contrast to a fully random sample, our sampling scheme contained no oversampling bias with regard to very recent content. Therefore, the generated sample was more representative of the whole observation period and accounted for the possibility that the nature of the tweets changed notably over time.

### 2.3 Annotation

After generating the sample, the selected tweets were annotated through the Crowdbreaks platform^1^ [36], which uses crowdsourcing to annotate social media data. The platform allows for the creation of a question sequence that is then submitted in combination with a tweet as a task to Amazon Turk (MTurk)^2^. The question sequence contained three questions for each task: The first question was on the relevance of the tweet to the topic of CRISPR-Cas9, allowing “relevant” and “not relevant” as possible answers. The second question was on the sentiment (“positive”, “negative”, or “neutral”), and the third question was on the organism (“humans”,”human embryos”,“animals (other than human)”,“plants”,“bacteria”,“multiple”,“not specified”).

Before submitting the task to MTurk, the availability of the tweet was automatically checked. This was done in order to respect the user’s right to either delete their content or set it to private after the time of data collection. Filtering by tweets which were still available yielded the sample *S*_1_ (*n* = 22 513), which was subsequently annotated. In order to reach a consensus, each tweet was annotated by multiple annotators on multiple questions, resulting in an annotation data set *A*_0_ of 226 670 individual answers on 22 492 unique tweets.

On both the question of relevance as well as the sentiment, the resulting Fleiss’ kappa agreement scores [37] were 0.81 and 0.28, respectively (0 denotes chance agreement, and 1 denotes perfect annotator agreement). In order to detect workers with questionable performance, the annotators’ raw agreement was calculated, which denotes the fraction of the number of actual agreements over the number of possible agreements an annotator had with other annotators. An annotator was considered an outlier if this value was smaller than three standard deviations from the mean, the annotator had at least 20 possible agreements with other annotators, and was involved in at least 3 separate tasks. All annotations by outlier annotators were subsequently removed. Annotations of tweets for which a unanimous consensus of at least 3 independent annotators could be found were then merged into a dataset *A*_1_ (*n* = 17 090) containing one label per tweet for each of the 3 questions. Dataset *A*1 was used as a training dataset for the relevance classifier. Annotations belonging to tweets which were labelled as relevant were then exported into dataset *A*_2_ (*n*=16 822). Dataset *A*_2_ was then used as a training dataset for all other trained classifiers.

### 2.4 Training

In order to classify the data with regard to relevance, sentiment and organism, we constructed three classifiers, *M*_*R*_, *M*_*S*_ and *M*_*O*_, respectively. The classifiers tried to predict the respective labels from the text of the tweet alone. In the process, we analyzed the performance of four different classifier models: Bag of Words, Sent2Vec sentence embeddings [38] coupled with Support Vector Machines (SVMs) [39], FastText [40], and BERT [34]. The tokenization and word character encoding process was different for each model class. In order to evaluate the models, the cleaned annotation data was shuffled and split into a training (80%, *n*=4250) and test set (20%, *n*=1063). After an extensive model selection process, a fine-tuned version of BERT-large was selected as the best performing sentiment model with a macro-averaged F1-score of 0.727 (*F*1_positive_ = 0.827, *F*1_neutral_ = 0.715, *F*1_negative_ = 0.639). BERT was also found to be the best performing model for the relevance and organism classifiers, resulting in a macro-averaged F1-score of 0.91 (*F*1_related_ = 0.997, *F*1_unrelated_ = 0.823) and 0.89 (*F*1_humans_ = 0.873, *F*1_embryos_ = 0.762, *F*1_animals_ = 1, *F*1_plants_ = 0.889, *F*1_bacteria_ = 0.909, *F*1_not specified_ = 0.902).

### 2.5 Prediction

For the analysis, the best performing model (BERT) for relevance *M*_*R*_ was used to predict dataset *D*_1_ and yield the predicted dataset *P*_0_ (*n*=1 334 114) of same length containing a label for relevance. Next, all tweets predicted as not relevant were removed from *P*_0_, yielding dataset *P*_1_ (*n*=1 311 544). This dataset was then used to predict sentiment and organism using the models *M*_*S*_ and *M*_*O*_, resulting in the final dataset *P*_2_.

### 2.6 Analysis

In our analysis, we used the sentiments in relation to tweet activity (number of tweets), topics of the tweets (hashtags), organisms the tweets were talking about (predicted), and themes identified from previous studies on CRISPR mentioned earlier (through regular expressions), to gain different kinds of insights. Wherever we used sentiments for numerical calculations, we used +1 for positive, 0 for neutral, and −1 for negative sentiment. Further, we extrapolated the numbers for 2019 where applicable for better comparison since we only had data until May 31, 2019. The different parts of the analysis are explained below in more detail.

The first part of the analysis is concerned with the development of the sentiment in relation to the number of tweets over time. The detection of temporary deviation from the general sentiment was of particular interest. While we included all tweets for the analysis of activity, we excluded tweets with neutral sentiment for the analysis of sentiment to make deviations more visible. We aggregated activity and sentiments on a daily basis. For the sentiments however, the sentiment value of a specific day was determined by taking the mean value of all positive and negative sentiments within a sliding seven day window centered around that day (± 3 days). We then used scipy’s module for peak detection [41] in order to detect events of interest, using a relative prominence cut-off of 0.2. In order to identify potential sources for the change in sentiment, we manually identified major events that relate to CRISPR.

In the second part, we used the predictions of the model *M*_*O*_ and the sentiments to compare the development of the sentiment for different organisms. We calculated the mean sentiments over a month and excluded all months that did not have at least 100 tweets for the respective organism.

Third, we analyzed hashtags as a proxy for the topics a user was talking about in his or her tweet. The hashtag “CRISPR” was excluded from the analysis since CRISPR was the overarching topic all tweets had in common. We counted the occurrences of every hashtag per year. We used the exact hashtags and did not group similar hashtags. For example, the hashtags “crisprbaby” and “crisprbabies” were treated as different hashtags. We did this due to the difficulty of automatically matching similar hashtags, since they can be a composition of multiple words that made strategies like stemming not straightforward. For each hashtag and year, we then calculated the mean sentiment and selected the 15 most common hashtags for each year for further analysis. We then manually compared how these top 15 topics per year increased and decreased in popularity throughout the years, as well as how the sentiments for these topics changed.

In the fourth and last part of our analysis, we based our analysis on the earlier conducted studies. We conducted a literature search in scientific databases according to a predefined search strategy (see SI text C). The search was conducted in the Fall of 2017. We reviewed the resulting studies and identified the themes where people had a positive or negative attitude towards CRISPR, or that concerned them. Additionally, we added themes based on publications and events that occurred between Fall of 2017 and Summer 2019. In order to see if these themes were also present in the tweets, we derived regular expressions representing the themes (see Table A.2 for the themes and the regular expressions derived from them). The regular expressions then allowed to automatically check for matches on the entire data set as a proxy for the presence of a certain theme. The results of the analysis are presented in the next section.

## 3 Results

Figure 2 shows a temporal analysis of the predicted sentiments in relation to key historical events surrounding CRISPR. A sentiment of zero indicates an equal portion of positive and negative tweets, and the values 1 and −1 indicate a signal with only positive or negative tweets, respectively. Figure 2A shows the sentiment *s* between July 2015 and June 2019. The time period before July 2015 was excluded, as activity was too low for a high-resolution sentiment signal. The sentiment remains mostly positive with an average of 85% positive tweets and only 15% negative tweets. Especially over an initial time period until March 2017, the sentiment shows little variation. After that, the sentiment reveals a series of sharp negative spikes, on two occasions dropping below zero. Over the observed time period, the sentiment shows a slight negative trend (slope of −0.06 y^−1^), as indicated by the linear trend line in orange.

**Figure 2:**
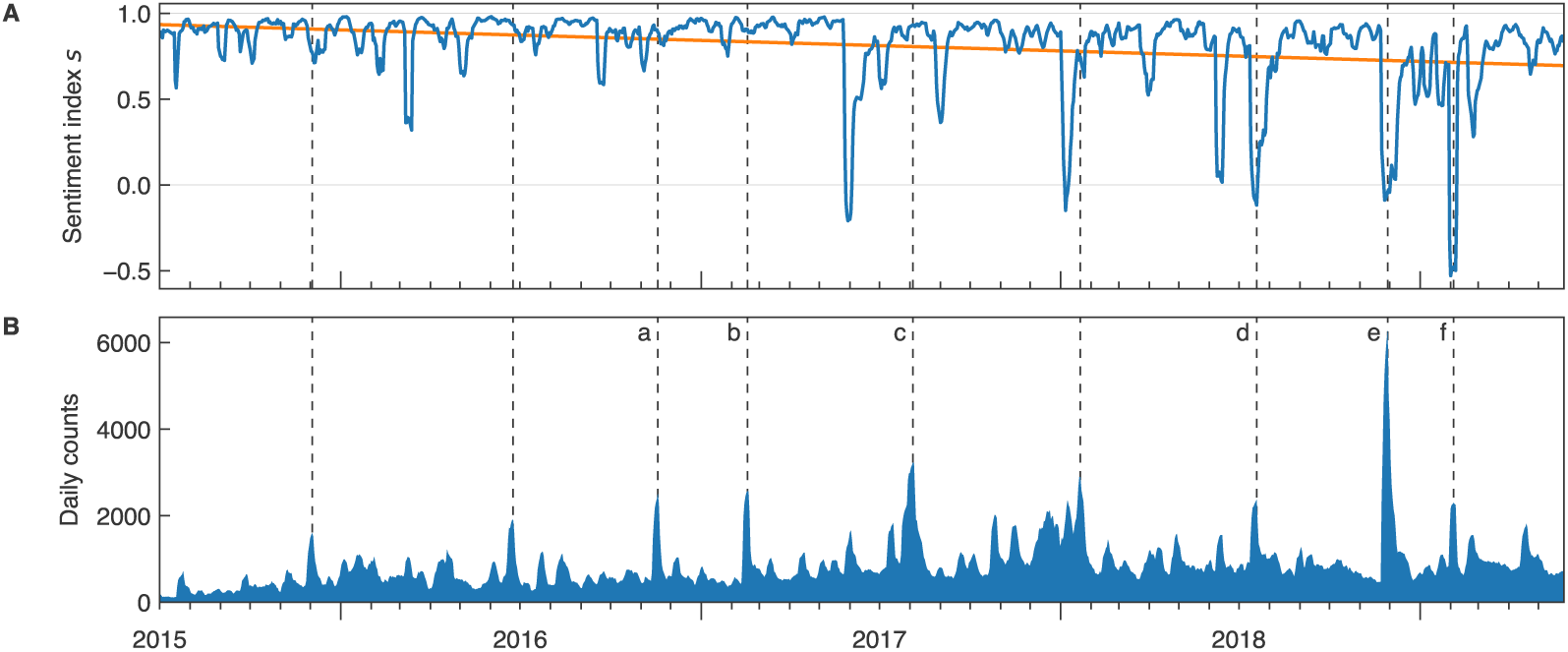
**A**) Predicted sentiment towards CRISPR between July 2015 to June 2019. The blue curve denotes the sentiment *s*, which is calculated as the mean of the weighted counts of positive and negative tweets over a centered rolling window of 7 days. The orange curve denotes a linear fit of the sentiment *s*. Up until May 2017, the sentiment towards CRISPR was mostly positive with minor variations. After that, the signal included multiple peaks of negative sentiment, at times with more negative than positive tweets. Overall, the orange line shows a slight negative trend, with a slope −0.06 y^−1^. **B**) Daily counts of all analyzed tweets. The blue area shows the daily sum of positive, negative and neutral tweets as the mean within a 7-day centered rolling window. All peaks above a relative prominence of 0.2 are marked with dashed lines. The peaks **a**-**f** denote peaks which coincide with certain events, which are referenced throughout the main text. The controversy commonly referred to as “CRISPR babies” marks the event associated with the highest average daily counts and a strong negative sentiment response (marked as peak **e**).

We then compared the sentiment curve to the observed activity surrounding CRISPR in the same time span, as shown in Figure 2B. Shown are the mean daily counts of the sample *P*_1_ over a sliding window of 7 days. Activity varies considerably with an average baseline of 1000 tweets per day and peaks of up to 6000 tweets per day.

We detected eight peaks of interest. They are marked with dashed lines in Figure 2B. When comparing peaks of high activity to the sentiment, it can be seen that peaks of high activity before mid 2018 did not result in a negative sentiment response. Peaks of strong negative sentiment started to appear in 2017 but it was not accompanied by the same level of activity until after 2018.

In a second step, major news events were manually mapped to coinciding peaks (for a full list see Table A.3). A subset of these peaks were marked with letters a-f in Figure 2, for illustrative purposes. In all cases the most retweeted tweet within days of the peak was linking a news article describing the event. The events include the first use of CRISPR in humans by a group of Chinese scientists in November 2016 (peak a), and the US Patent Office deciding in favor of the Broad Institute (peak b). Both of these events did not lead to a significant change in sentiment. Peak c coincides with the publication of a study which reported the correction of a mutation in human embryos [42], causing widespread media attention and, as before, did not cause a drop in sentiment. However, in July 2018, a study by the Wellcome Sanger Institute warned about serious side effects, such as cancer, which CRISPR could have when used in humans [43] (peak d). This peak led to a clear negative response in the sentiment index, and marks the first negative peak with high media attention. When researcher He Jiankui revealed to have created the world’s first genetically edited babies in November 2018 [16] (peak e), the highest activity was recorded. Although He’s revelation caused a strong negative signal, the strongest negative sentiment was recorded shortly after in February 2019 (peak f). This event coincides with the re-emergence of a news story from August 2017 when biohackers managed to encode a malware program into a strand of DNA [44].

In order to improve our understanding of the sentiment signal, the data was predicted with respect to which organism each tweet was about (see methods section 2.6). All relevant data (*n*=1 311 544) was predicted by organism into the classes animals (7.6%), bacteria (2.4%), embryos (4.3%), humans (3.0%), plants (4.9%) and not specified (50.6%). As expected, more than half of all tweets do not specifically refer to an organism in context with CRISPR. Animals (e.g. mice for animal testing) are the second largest group, whereas human and embryos combined make up for the third largest group. Tweets specifically mentioning CRISPR in the context of bacteria were rather rare.

Figure 3A shows the monthly sentiment for each organism class, which are based on the monthly counts shown in Figure 3B. Out of all classes, embryos exhibited the most negative leaning sentiment (with a mean sentiment *μ*_*s*_ of 0.13). Embryos was also the class with the strongest variations of sentiment based on the monthly standard deviation (*σ*_*s*_ =0.28). A relatively high sentiment was measured for the classes animals (*μ*_*s*_ =0.70), bacteria (*μ*_*s*_ = 0.64), and plants (*μ*_*s*_ = 0.61). For the class humans, the average sentiment was relatively high (*μ*_*s*_ =0.58) but showed a clear dip in sentiment in the months following November 2018. The class unspecified showed a slightly lower mean of 0.45 compared to other classes.

**Figure 3:**
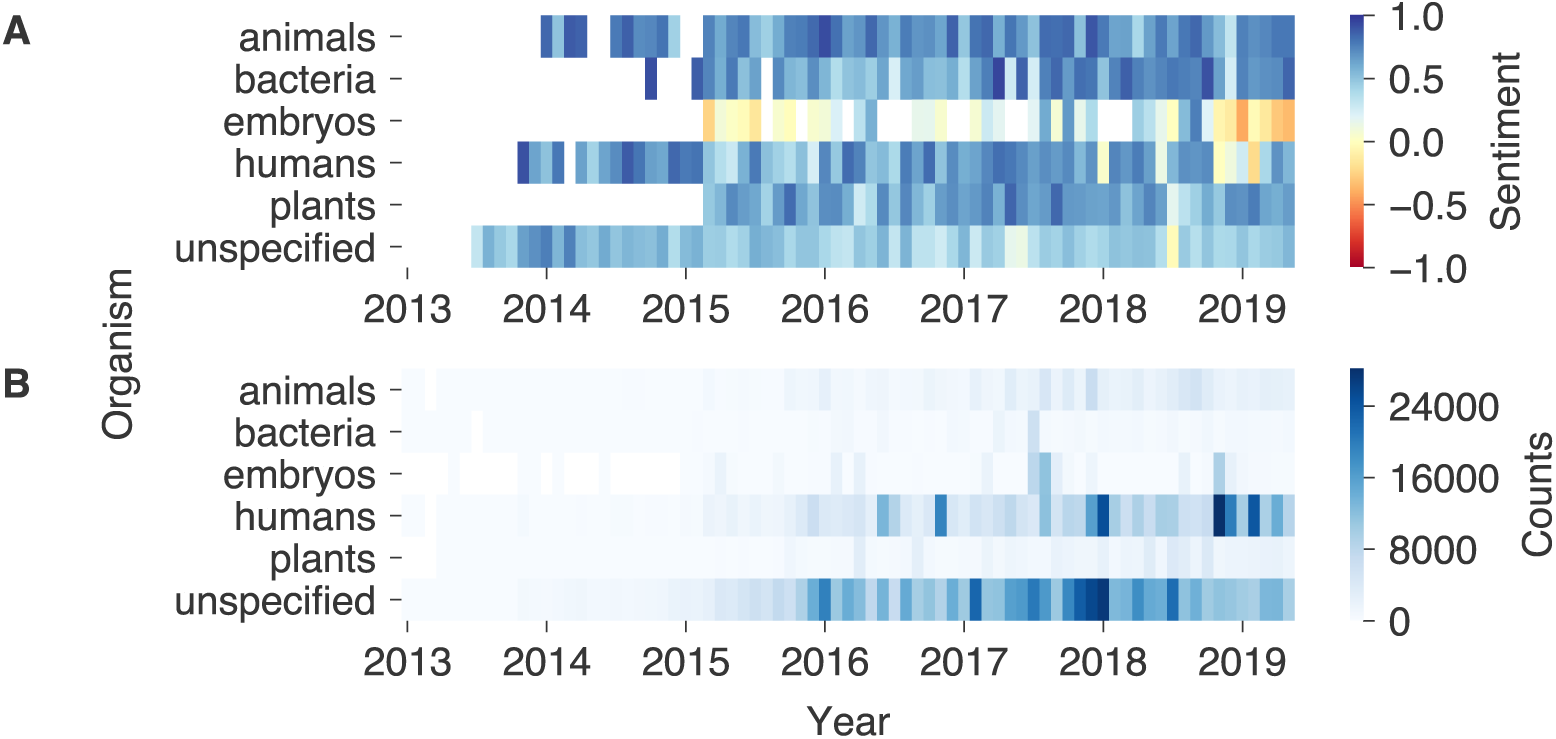
**A**) Heatmap of monthly sentiments by predicted organism. The sentiments were calculated as the mean of the weighted counts by sentiment (the weights include −1, 0 and 1 for negative, neutral and positive tweets) for each month and organism class. Blue and red colors indicate positive and negative sentiment values, respectively. The sentiments of heatmap cells with less than 100 tweets of that month and organism were left transparent. Tweets classified as relating to animals, bacteria and plants showed mostly a positive sentiment throughout the observed timespan. For the human category the sentiment was positive up until November 2018, at which it began to drop. The sentiment with respect to embryos was much lower and further declined after November 2018. **B**) Monthly counts by predicted organism. The counts serve as the basis for the sentiment calculated in panel A. The monthly counts increased throughout the years for all organisms. A majority of tweets were of class unspecified. For the classes “human” and “embryo”, occasional months of high activity was observed.

The most frequent hashtags of every year revealed the topics of highest interest and how they evolved over time (see Figure 4). Naturally, the occurrences of individual hashtags increased over the years along with the total number of tweets. Certain very common hashtags such as #dna, #science, #biotech, or #geneediting and #genomeediting, appeared as top hashtags in multiple years. When relating the hashtags with the sentiment of the text they appeared in, we can see that most of these common hashtags were used in the context of a positive or very positive sentiment. The three hashtags with the most positive sentiments and more than 100 occurrences wer #cancer with a mean sentiment of 0.85 in 2015, #hiv with 0.90 in 2016, and #researchhighlight with 1.0 in 2019. It is also notable that #science was among the five most common hashtags in every year except for 2013, and was consistently related to a positive sentiment between 0.52 and 0.74.

**Figure 4:**
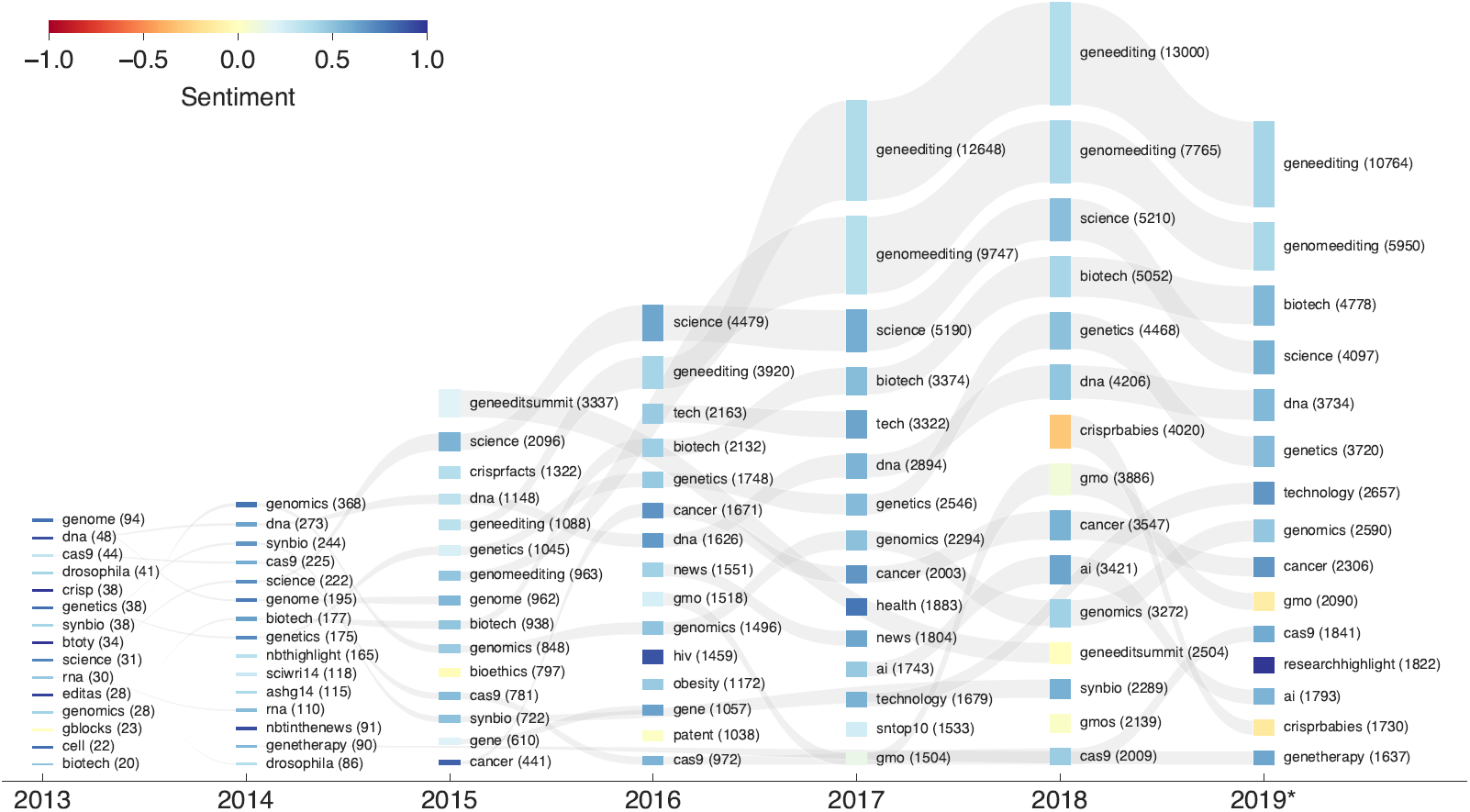
Visualization of the sentiment associated with the most frequent hashtags every year. For every year, the 15 hashtags with the highest counts for that year are included (the hashtag #crispr was excluded). The hashtags are sorted by yearly counts (indicated by the bar height) where the hashtag with the highest count is at the top. The color represents the average sentiment for the respective hashtag, blue color representing a very positive, and red color representing a very negative sentiment. If a hashtag is listed in multiple years, the occurrences are linked with a gray band. The number of tweets with the hashtag is indicated in parentheses next to the respective hashtag. For the year 2019, the counts were extrapolated from the months until June to the full year.

Only a few hashtags were related to negative sentiments (#crisprbabies in 2018 and 2019, #gmo in 2019, #bioethics in 2015, and #geneeditsummit in 2018). The most prominent was #crisprbabies with a mean sentiment of −0.30 for 2018, and −0.13 in 2019 (see Table A.4 for a full list of the counts and sentiments of the most used hashtags by year). It is worth noting that the hashtag #geneeditsummit only appeared in 2015 and in 2018, and that its associated sentiment dropped from 0.20 to −0.01. The hashtag refers to the two summits on Human Genome Editing, which were held in Washington D.C. in 2015 and in Hong Kong in November 2018, coinciding with the first gene editing of viable human embryos. Similarly, the hashtag #gmo became slightly more negative in 2018, with a mean sentiment of 0.09 compared to the years before with 0.24 in 2016, and 0.14 in 2017, and even dropped to −0.11 in 2019. The hashtag #bioethics only appeared in 2015 and was associated with a relatively low sentiment of −0.02. This may highlight the various ethical concerns raised during the 2015 Human Genome summit.

In comparison to the hashtags, the themes derived from previous studies can relate the Twitter discussion to known themes of interest to the public (see methods section 2.6 for a description of the analysis). The six themes that were matched most are presented in Figure 5 and grouped by positive, neutral, and negative sentiments. The themes include “genome” (with a total count of 526 612), “baby” (68 269), “disease” (64 180), “embryo” (49 085), “treatment” (35 864), and “mutation” (34 884). Unsurprisingly, the theme “genome” was matched most frequently, occurring in 35% of all tweets. Themes have distinct occurrence patterns for each sentiment and reveal spikes in certain years. The most significant change in occurrences happened for the theme “baby” which increased substantially from 2017 to 2018, likely associated with the “CRISPR babies” scandal in November 2018. While a spike could be observed for all three sentiments, the increase was far more pronounced in the neutral and negative class (panels 5A and 5B). The theme “mutation” was clearly negatively connotated, showing a negative peak in 2017 when risks about potential side effects of CRISPR surfaced. Relative to other themes, the themes “disease” and “treatment” were major themes in a discussion correlated with a positive sentiment.

**Figure 5:**
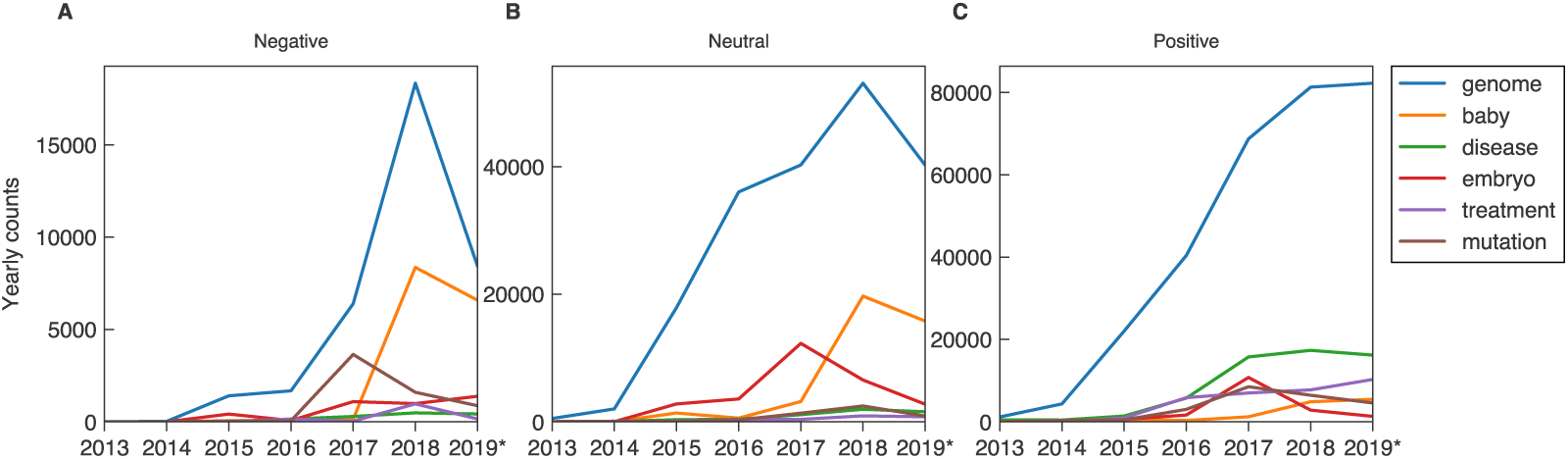
Yearly occurrences of themes. Multiple themes with distinct regex patterns were matched to the text of tweets, and the six most frequent themes were selected. Panels A, B, and C show the yearly counts of themes when grouped by negative, neutral and positive sentiment, respectively. The reported themes show distinct occurrence patterns depending on sentiment, yielding an aggregated picture of the discussion surrounding CRISPR throughout the years. Most counts increase over the years as the number of tweets increases overall. For the year 2019, the counts were extrapolated from the months until June to a full year.

## 4 Discussion

We have generated the first high-resolution temporal signal for sentiments towards CRISPR on Twitter, spanning a duration of more than 6 years. Our results suggest that, overall, the CRISPR technology was discussed in a positive light, which aligns well with a previous study which considered the coverage of CRISPR in the press [33]. However, more recently the sentiment reveals a series of strong negative dips, pointing to a more critical view. The frequency and magnitude of these dips has increased since 2017, which is underlined by the overall declining sentiment. It is noteworthy that the dips usually coincide with high activity, suggesting that most people are only exposed to the topic of CRISPR when it is presented in an unfavorable way.

The matching of activity peaks with events further ties the aforementioned dips to real world events. This suggests that the changes in the sentiment don’t just occur as result of a self-contained discussion but also reflect people’s conversations held outside Twitter. Even more so, the peak detection potentially allows to timely identify significant incidents that can shape public discourse and opinion. In addition, our results indicate a connection between the content of the conversation on and off Twitter.

As shown in the breakdown of sentiment by organism, the negative sentiment was stronger in the embryo and human class, but stayed mostly positive towards other organisms. The data therefore reflects the many ethical issues related to human germline editing. However, criticism may not be targeted at the use of CRISPR in humans per se: Hashtags such as #hiv or #genetherapy were connected to very positive sentiments, which suggests a positive attitude towards developing CRISPR for the use in medical treatment. This aspect is further confirmed when considering the sentiment of themes such as “treatment” or “disease”. These observations are in line with several surveys in which participants demonstrated a strong support of CRISPR for the use in medical treatment, but were critical regarding modifications of human germline cells [28, 29, 30, 31, 32].

The dataset which includes continuous observations over a long period of time allows to draw conclusions about the public perception of CRISPR both on short and long time scales. For example, when the article on biohacking re-emerged in 2019 (peak f), shortly after the discussions around CRISPR babies, it was discussed in significantly more negative terms than at the time of its publication in 2017. Therefore, the intermediate developments must have had a negative influence on the perception of the event. This is in line with the overall negative trend. The influence of single events on the overall discussion also manifests itself in the presence and absence of themes in the discussion. While the theme “mutation” was discussed intensely in 2017, its occurrence in tweets dropped again drastically in the following year in which “baby” became the single most occuring theme by a large margin.

Our results support the use of Twitter and similar platforms for the study of public discourse. The discussion about a subject matter can be investigated in real-time, in depth on the level of individual statements, and on the basis of existing data. The insights gained through such studies can bring new issues to light, indicate which topics need extra attention with respect to ethical considerations and policy making, and allow to respond quicker to technological advancements. In addition the presented method offers a novel approach to promote public engagement, especially in the area of biotechnologies and health care, as argued for by the Nuffield Council on Bioethics [45].

### 4.1 Limitations

Although the predicted sentiment index seems to overlap well with survey results, it cannot be directly used as a substitute for an opinion poll. Polling allows for the collection of answers to specific questions of interest instead of inferring them from public statements. Furthermore, the Twitter community is not necessarily representative of the whole population of a country. However, sentiment analysis avoids downsides of traditional methods such as response bias, and provides more detailed insights through access to fine grained data of the online discussion.

We acknowledge that most people’s opinions might not fit into the positive, neutral and negative classes presented in this study. We therefore tried to counteract this problem by not only categorizing the data by sentiment, but also by relevance and organism, allowing for a better understanding of the measured sentiment. Furthermore, we recognize the challenging nature of deducing someone’s true opinion based on a short message alone and the fact that it is only possible within a statistical margin of error. We believe however that by employing extended preprocessing, filtering, and state of the art machine learning, we can capture certain trends on a larger scale.

### 4.2 Conclusions and future direction

We have demonstrated that the sentiment analysis of tweets provides a high resolution picture of the ongoing debate on CRISPR, allowing us to study the evolution of the discourse while extending the capacity of traditional methods. Further, the presence of the same themes that have been identified in existing studies confirms the validity of our signal with respect to content. The existence of events that match the activity peaks also indicates the sensitivity of the signal towards off-Twitter incidents. Therefore, our approach offers an additional method to surveys and which can be deployed to get richer information, higher sample size and higher temporal resolution.

Future work can go beyond the deduction of sentiments and shed more light on the nature of discussions and arguments raised and how they influence each other, giving a better idea of the reasoning behind people’s opinions. Furthermore, specific topics, such as the discussion surrounding a potential moratorium of CRISPR, may be analyzed in more detail and provide actionable outcomes.

Since the presented analysis can automatically process a large amount of data in almost real-time, it extends the traditional toolset of empirical methods for discourse analysis. It may therefore help analyzing public opinion and support policy and decision making.

## 5 Data Availability Statement

All data and code for this analysis can be found in our public repository: https://gitlab.ethz.ch/digitalbioethics/crispr-sentiment-analysis.

## 6 Acknowledgements

We thank Agata Ferretti for the support during the initial literature review and Ellen Lapper for proofreading.

## A Appendix

### A.1 Yearly counts

**Table A.1:** Number of tweets per year since January 1, 2013, until May 31, 2019. A steady increase in volume can be observed. For the year 2019 the number was extrapolated from 163 553 until May 31, 2019.

| Year | Number of tweets |
| --- | --- |
| 2013 | 4818 |
| 2014 | 20 003 |
| 2015 | 131 268 |
| 2016 | 304 751 |
| 2017 | 437 928 |
| 2018 | 445 723 |
| 2019 | 392 527 |
| **Total** | **1 508 044** |

### A.2 Model performance

**Figure A.1:**
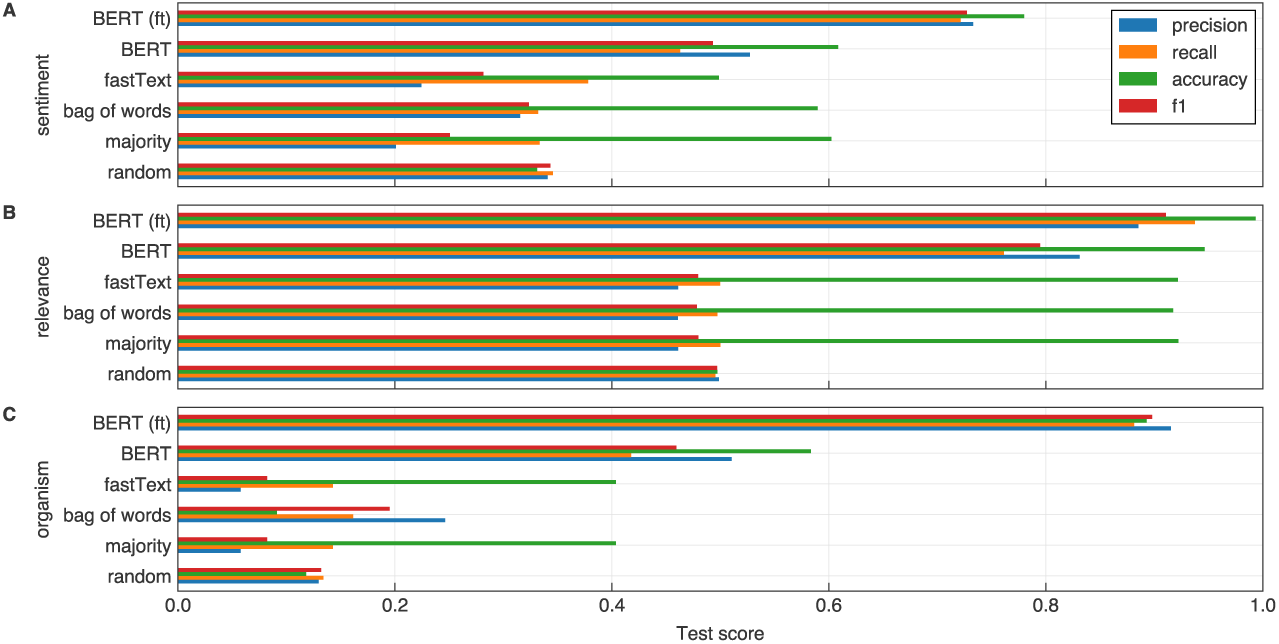
Classification scores for different models. Subfigures **A, B** and **C** correspond to three different classifiers trained for sentiment, relevance and organism, respectively. They y-axis shows best corresponding model after hyperparameter search was performed for a specific model type. The model types are random (pick a class at random), majority (always pick the most frequent class), bag of words, fastText, BERT and a fine-tuned version of BERT-large (denoted as BERT ft). The x-axis denotes the test performance scores of accuracy (green), and macro-averaged precision (blue), recall (orange) and F1 scores (red). The fine-tuned BERT model was the best performing model for all three classification problems irrespective of the metric used.

### A.3 Preliminary literature review search strategy and databases

Databases used: PubMed, Scopus, Web of science. Matching query in articles’ title only: `((crispr OR gene-editing OR “genome editing”) AND (attitudes OR opinions OR perspectives OR believes OR reactions OR public))` 103 publications were identified by the search (24 PubMed, 41 Scopus, 38 Web of Science). A total of 4 articles were included in the full-text analysis after duplicate removal and exclusion through abstract screening based on exclusion criteria:

- The article is not focussing on CRISPR
- The article is not referring to human subjects
- The article is not considering public opinions/attitudes
- The article is not an empirical study

#### Resulting documents

- Blendon, R. J., Gorski, M. T., & Benson, J. M. (2016). The public and the gene-editing revolution. New England Journal of Medicine, 374(15), 1406-1411.
- McCaughey, T., Sanfilippo, P. G., Gooden, G. E., Budden, D. M., Fan, L., Fenwick, E., … & Liang, H. H. (2016). A global social media survey of attitudes to human genome editing. Cell stem cell, 18(5), 569-572.
- Scheufele, D. A., Xenos, M. A., Howell, E. L., Rose, K. M., Brossard, D., & Hardy, B. W. (2017). US attitudes on human genome editing. Science, 357(6351), 553-554.
- Weisberg, S. M., Badgio, D., & Chatterjee, A. (2017). A CRISPR New World: Attitudes in the Public toward Innovations in Human Genetic Modification. Frontiers in Public Health, 5.

### A.4 Themes and regex patterns

**Table A.2:**
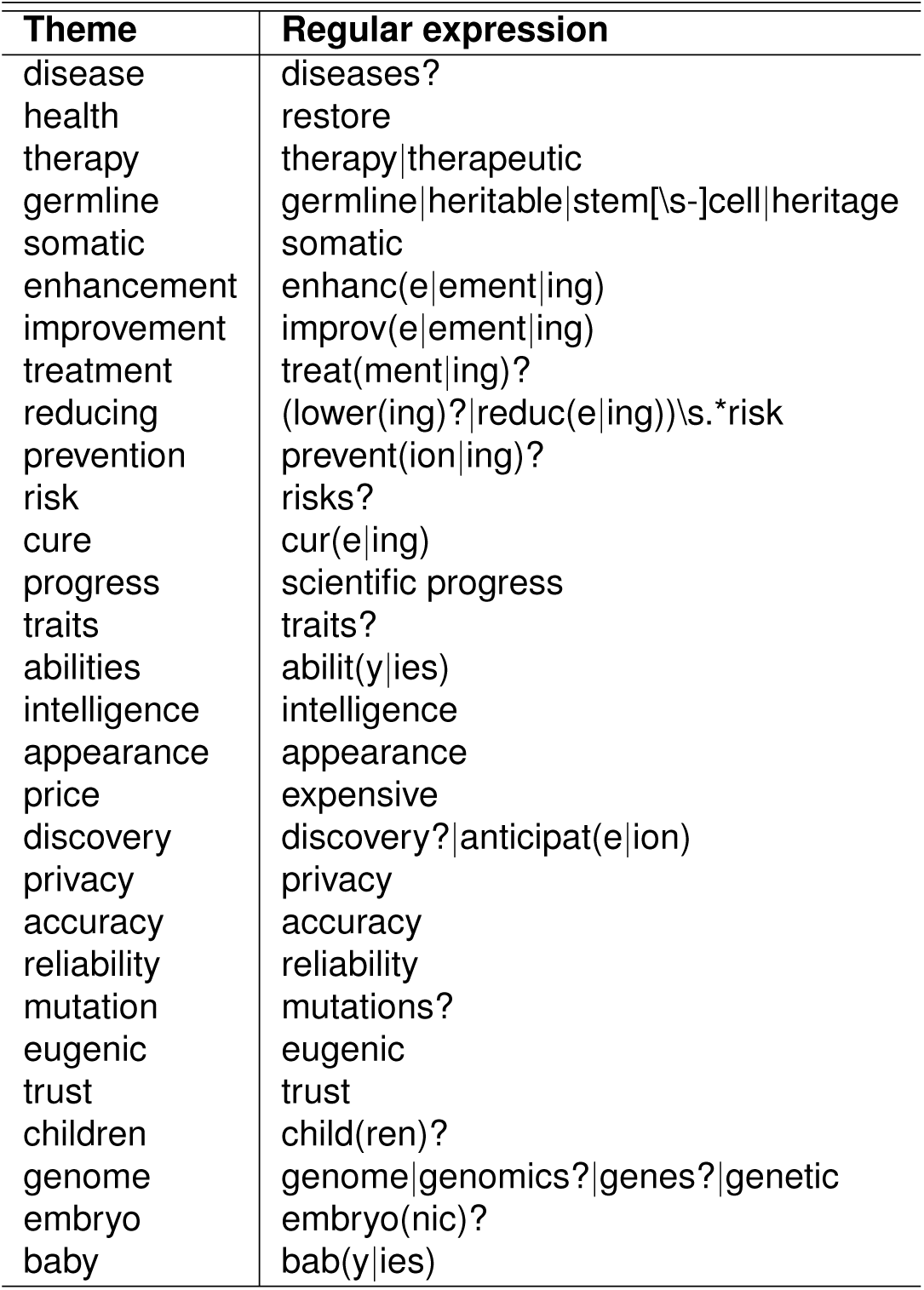
Derived themes and corresponding regex patterns from preliminary literature review.

### A.5 Identified events

**Table A.3:**
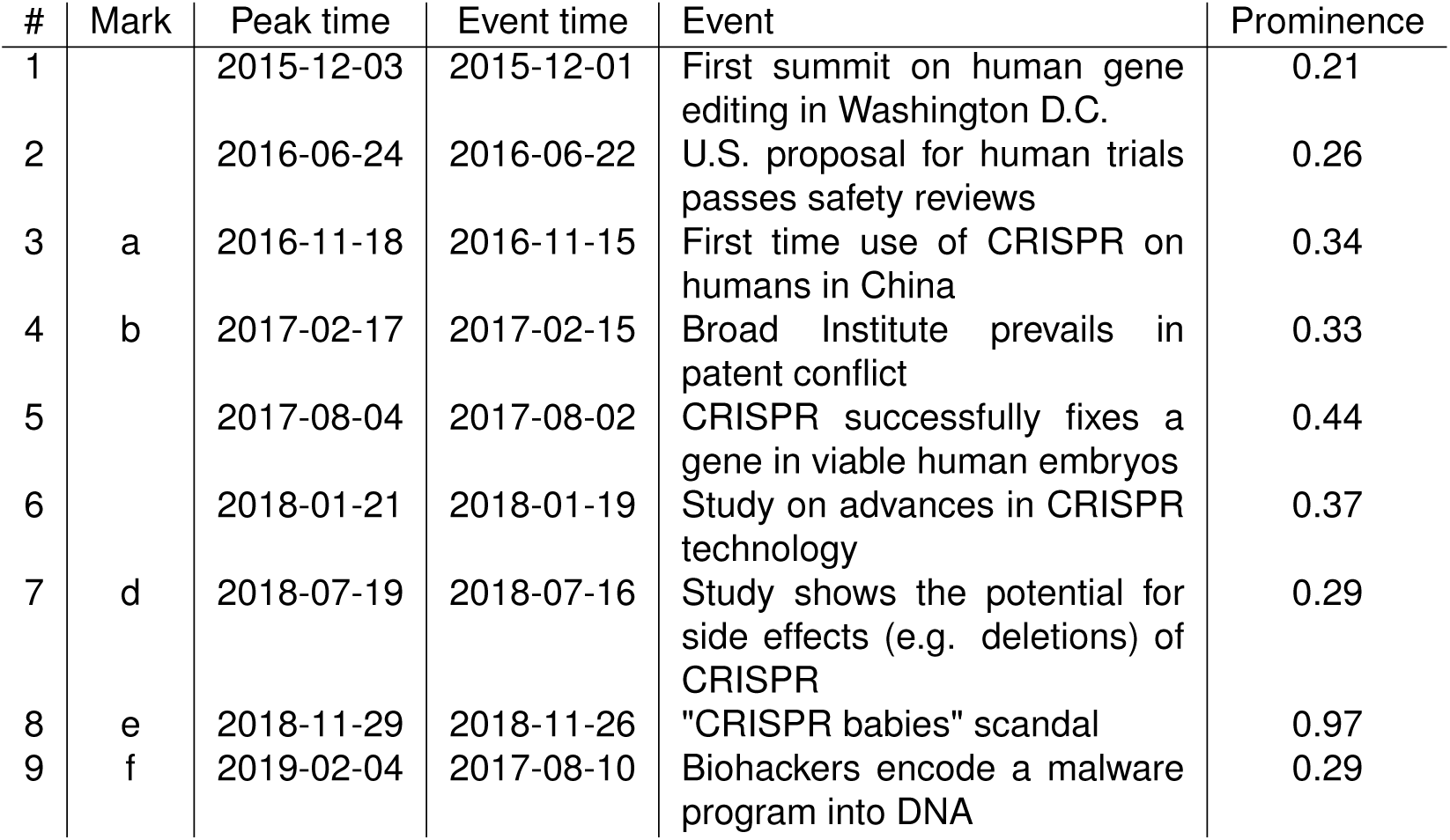
Selected events with a peak prominence above 0.2. The marks correspond to the selected events in Figure 2. Peak times have been automatically detected as described in the methods section. The corresponding events have been inferred from visual inspection of the data.

| # | Mark | Peak time | Event time | Event | Prominence |
| --- | --- | --- | --- | --- | --- |
| 1 |  | 2015-12-03 | 2015-12-01 | First summit on human gene editing in Washington D.C. | 0.21 |
| 2 |  | 2016-06-24 | 2016-06-22 | U.S. proposal for human trials passes safety reviews | 0.26 |
| 3 | a | 2016-11-18 | 2016-11-15 | First time use of CRISPR on humans in China | 0.34 |
| 4 | b | 2017-02-17 | 2017-02-15 | Broad Institute prevails in patent conflict | 0.33 |
| 5 |  | 2017-08-04 | 2017-08-02 | CRISPR successfully fixes a gene in viable human embryos | 0.44 |
| 6 |  | 2018-01-21 | 2018-01-19 | Study on advances in CRISPR technology | 0.37 |
| 7 | d | 2018-07-19 | 2018-07-16 | Study shows the potential for side effects (e.g. deletions) of CRISPR | 0.29 |
| 8 | e | 2018-11-29 | 2018-11-26 | "CRISPR babies" scandal | 0.97 |
| 9 | f | 2019-02-04 | 2017-08-10 | Biohackers encode a malware program into DNA | 0.29 |

### A.6 Top hashtags counts and sentiments

**Table A.4:**
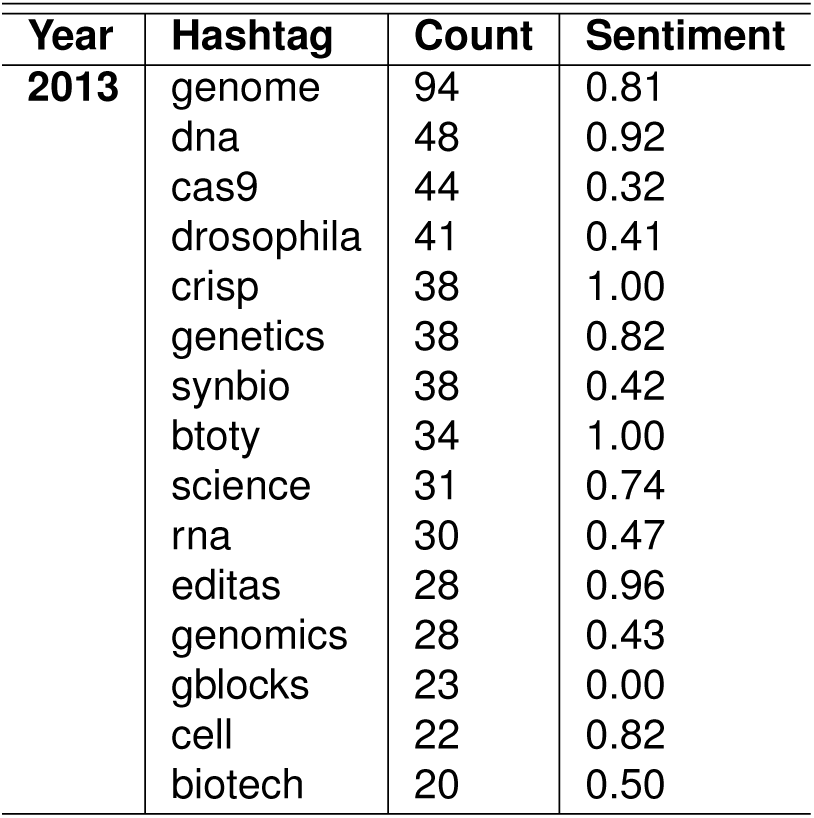
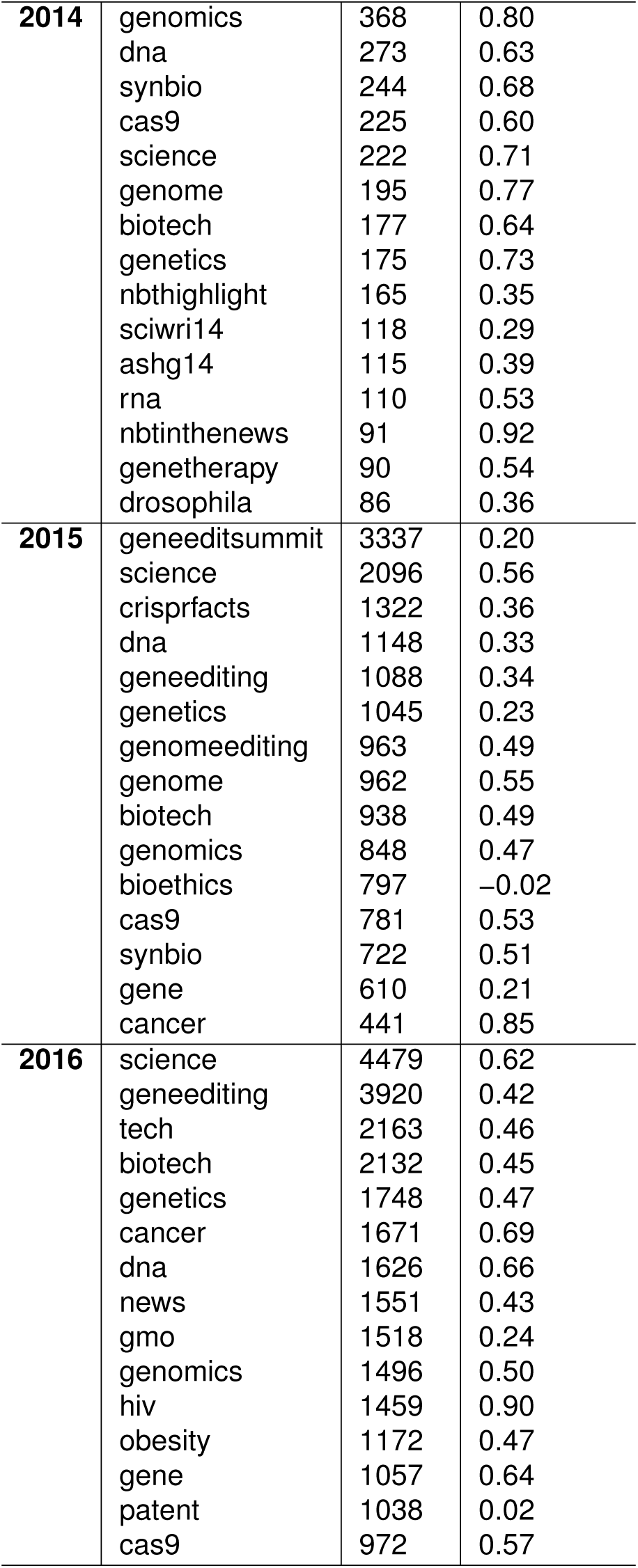
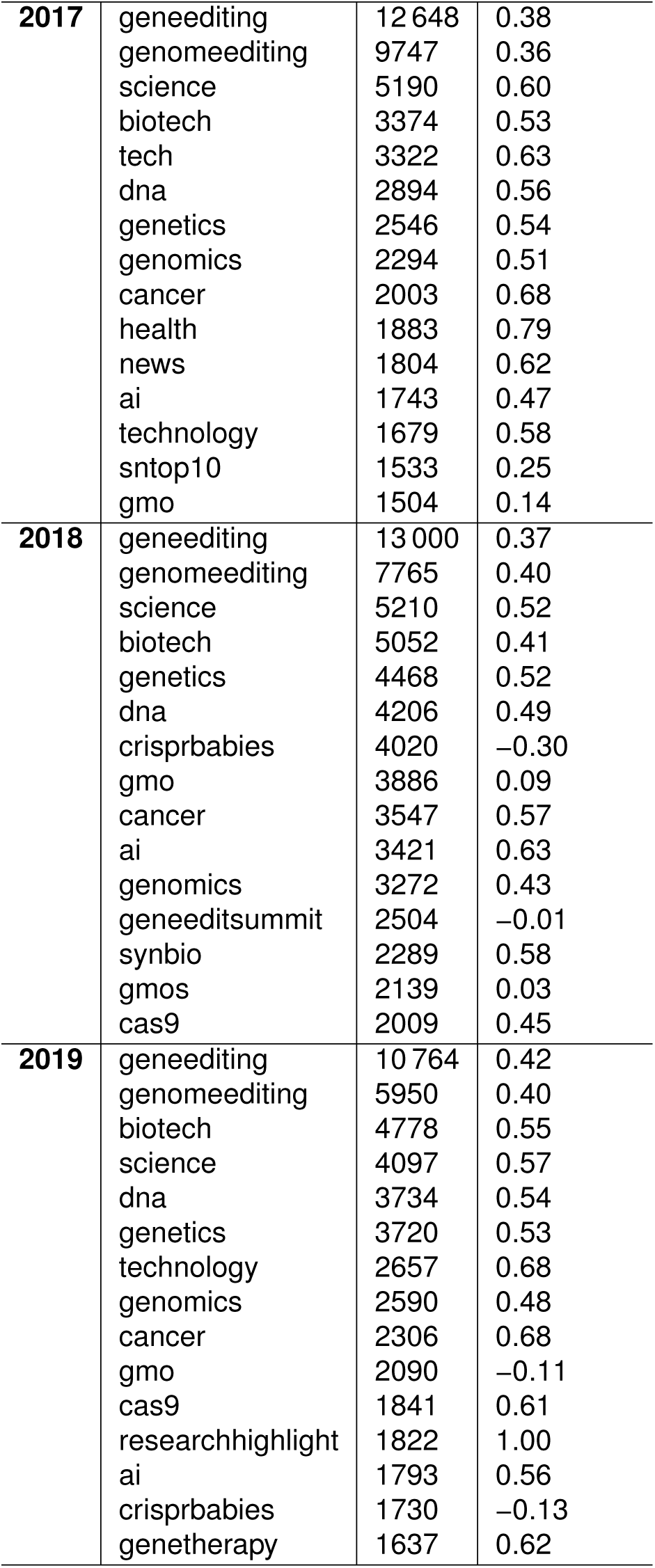
List of top 15 hashtags and corresponding counts and sentiments by year.

## Footnotes

1 https://www.crowdbreaks.org

2 https://www.mturk.com/

